# Monkeys are Curious about Counterfactual Outcomes

**DOI:** 10.1101/291708

**Authors:** Maya Zhe Wang, Benjamin Y. Hayden

## Abstract

While many non-human animals show basic exploratory behaviors, it remains unclear whether any animals possess human-like curiosity. We propose that human-like curiosity satisfies three formal criteria: (1) willingness to pay (or to sacrifice reward) to obtain information, (2) that the information provides no instrumental or strategic benefit (and the subject understands this), and (3) the amount the subject is willing to pay scales with the amount of information available. Although previous work, including our own, demonstrates that some animals will sacrifice juice rewards for information, that information normally predicts upcoming rewards and their ostensible curiosity may therefore be a byproduct of reinforcement processes. Here we get around this potential confound by showing that macaques sacrifice juice to obtain information about counterfactual outcomes (outcomes that could have occurred had the subject chosen differently). Moreover, willingness-to-pay scales with the information (Shannon entropy) offered by the counterfactual option. These results demonstrate human-like curiosity in non-human animals according to our strict criteria, which circumvent several confounds associated with less stringent criteria.

## INTRODUCTION

Curiosity is a major driver of exploration and learning. The term curiosity generally refers to information-seeking behavior that is intrinsically motivated (Golman & Loewenstein, 2016; Gottlieb, Oudeyer, Lopes, & Baranes, 2013; Kidd & Hayden, 2015; Loewenstein, 1994; Oudeyer, Kaplan, & Hafner, 2007). The intrinsic factor distinguishes curiosity from strategic forms of information seeking, such as exploration in bandit tasks (Daw, O’Doherty, Dayan, Seymour, & Dolan, 2006; Hayden & Platt, 2009). Thus, a stringent definition of curiosity refers to information seeking that reduces a decision-maker’s *information gap* without producing immediate or even potential reward or strategic benefits (Golman & Loewenstein, 2015; 2016). By this definition humans are curious (Berlyne, 1966; Gottlieb et al., 2013; Gruber, Gelman, & Ranganath, 2014; Kang et al., 2009; Loewenstein, 1994). For example, many people will pay money to for answers to trivia questions or to solve crossword puzzles even when those answers are patently useless (i.e. trivial,Kang et al., 2009; Gruber, Gelman, & Ranganath, 2014).

Thus we propose a conservative definition for human-like curiosity that requires (1) a willingness to pay for information, (2) the information is strategically useless, and (3) the information-seeking tendency increases with the amount of information provided (at least up to a point, see **Discussion**). Is it not clear whether non-human animals possess human-like curiosity according to this conservative definition (Kidd & Hayden, 2015). Many animals naturally explore their surroundings (Berlyne, 1966). For example, monkeys seek specific information while solving mechanical puzzles without immediate extrinsic motivations (Davis, Settlage, & Harlow, 1950; H. F. Harlow, 1950; H. F. Harlow, Harlow, & Meyer, 1950). Rats also show spontaneous exploration of unfamiliar maze sections without explicit reward or task (Dember, 1956; Hughes, 1968; Kivy, Earl, & Walker, 1956; Tolman, 1948). However, these information-seeking behaviors may be byproducts of natural foraging behavior where exploration or problem solving could lead to food (Emery & Clayton, 2004; Thorndike, 2017). Indeed, in these contexts, the animal may believe the actions performed could lead directly to reward (Menzel, 1991). Another practical limitation of these studies is the difficulty of formally quantifying the information gap, and thus showing that demand scales with information amount.

These problems have motivated scholars to focus on more controlled paradigms. Rigorous experiments have quantified information-seeking behavior under controlled conditions in species ranging from *Caenorhabditis elegans* worms (Calhoun, Chalasani, & Sharpee, 2014) to rhesus macaques (Averbeck, 2015; Costa, Monte, Lucas, Murray, & Averbeck, 2016; Noonan et al., 2010; Pearson, Hayden, Raghavachari, & Platt, 2009; Walton, Behrens, Buckley, Rudebeck, & Rushworth, 2010; Whittle, 1988). However, information in these tasks generally offers strategic benefits that could lead to greater future rewards. Likewise, in some uncertain contexts, animals prefer risky options; these options may be favored because they provide more information (Heilbronner & Hayden, 2013). Again, however, other non-curiosity-related factors may explain risky choice in these contexts, such as erroneous belief that stochastic processes are actually patterned (Blanchard, Wolfe, Vlaev, Winston, & Hayden, 2014; Hayden & Platt, 2007).

Another paradigm used to demonstrate curiosity has been the temporal resolution of un-certainty paradigm (Blanchard, Hayden, & Bromberg-Martin, 2015a; Bromberg-Martin & Hikosaka, 2009; Kidd, Palmeri, & Aslin, 2013). Many animals will sacrifice small amounts of food to obtain information about upcoming stochastic rewards (sometimes known as observing behavior,Blanchard et al., 2015a; Bromberg-Martin & Hikosaka, 2009; Bromberg-Martin, Matsumoto, & Hikosaka, 2010; Gipson, Alessandri, Miller, & Zentall, 2009; Roper, 1999; Stagner & Zentall, 2010; Zentall & Stagner, 2011). However, two major factors raise doubt about the ability of observing behavior tasks to demonstrate curiosity. First, Pavlovian learning may bias instrumental choice actions (Beierholm & Dayan, 2010; Dayan, Niv, Seymour, & Daw, 2006). That is, information in these tasks reliably predicts upcoming rewards. In a modified temporal difference learning model, informative cues increases the subjects’ engagement. The engagement maintains the representation of subjective value associated with the cue and the states associative with the objective reward. Thus, a non-informative (non-predictive) cue would lead to disengagement and a decrease in representation. This model predicts the preference for informative choice due not to the curiosity for information, but to the engaged and maintained representation of predicted value. Another related possibility is that animals may “superstitiously” believe that their choices with information could affect upcoming rewards (Vasconcelos, Monteiro, & Kacelnik, 2015).

Most of these problems stem from the direct association between information and immediate or potential rewards. Therefore, one way to avoid these confounds is to focus on curiosity about counterfactual outcomes. The term counterfactual refers to outcomes associated with options that were not chosen (the terms hypothetical and fictive are also sometimes used,Abe & Lee, 2011; Hayden, Pearson, & Platt, 2009). Monkeys can recognize counterfactual outcomes; their responses to counterfactual information indicate that understand its meaning, and do not simply respond as they would to conditioned reinforcers (Abe & Lee, 2011; Hayden et al., 2009; Rosati & Hare, 2013). Therefore, counterfactual outcomes can potentially help avoid some problems associated with paradigms in which curiosity-driving information relates to potential rewards.

We devised a *counterfactual information task* for rhesus macaques. On each trial, subjects chose between two gambles with independently generated stakes, probabilities, and counterfactual information status. That is, some options offered, if chosen, the promise of information about the result of the unchosen gamble; other options did not offer that information. We found that monkeys actively sought out information of counterfactual outcome, despite the lack of its instrumental benefits for current or future reward, and their preference for informative options scaled with the amount of information (i.e. Shannon entropy).

## RESULTS

### Monkeys Seek Counterfactual Information

Two rhesus macaques performed a novel gambling task designed to measure the subjective value placed on information about counterfactual outcomes (**Figure 1A** and **Methods**). On each trial of the *counterfactual information task*, monkeys chose between two gambles (offer 1 and offer 2) presented on the left and the right side of the screen. Presentation was asynchronous; order of sides was randomized. Gambles differed in three dimensions: *payoff* (small: 125 uL of water; medium: 165 uL; large: 250 uL, indicated by yellow, blue, or green color); *probability* (0 to 1 in 0.01 increments, indicated by bar section height); and *informativeness* (that is, whether the choice would lead to counterfactual information, indicated by the presence of an inscribed circle, **Figure 1A**). Payoff, probability, and informativeness were independently randomized for each offer on each trial. On 50% of the trials, only one option was informative (*info choice* trials), on 25% of the trials, both options were informative (*forced info* trials), and on the remaining 25%, neither option was informative (*no info* trials).

**Figure 1.**
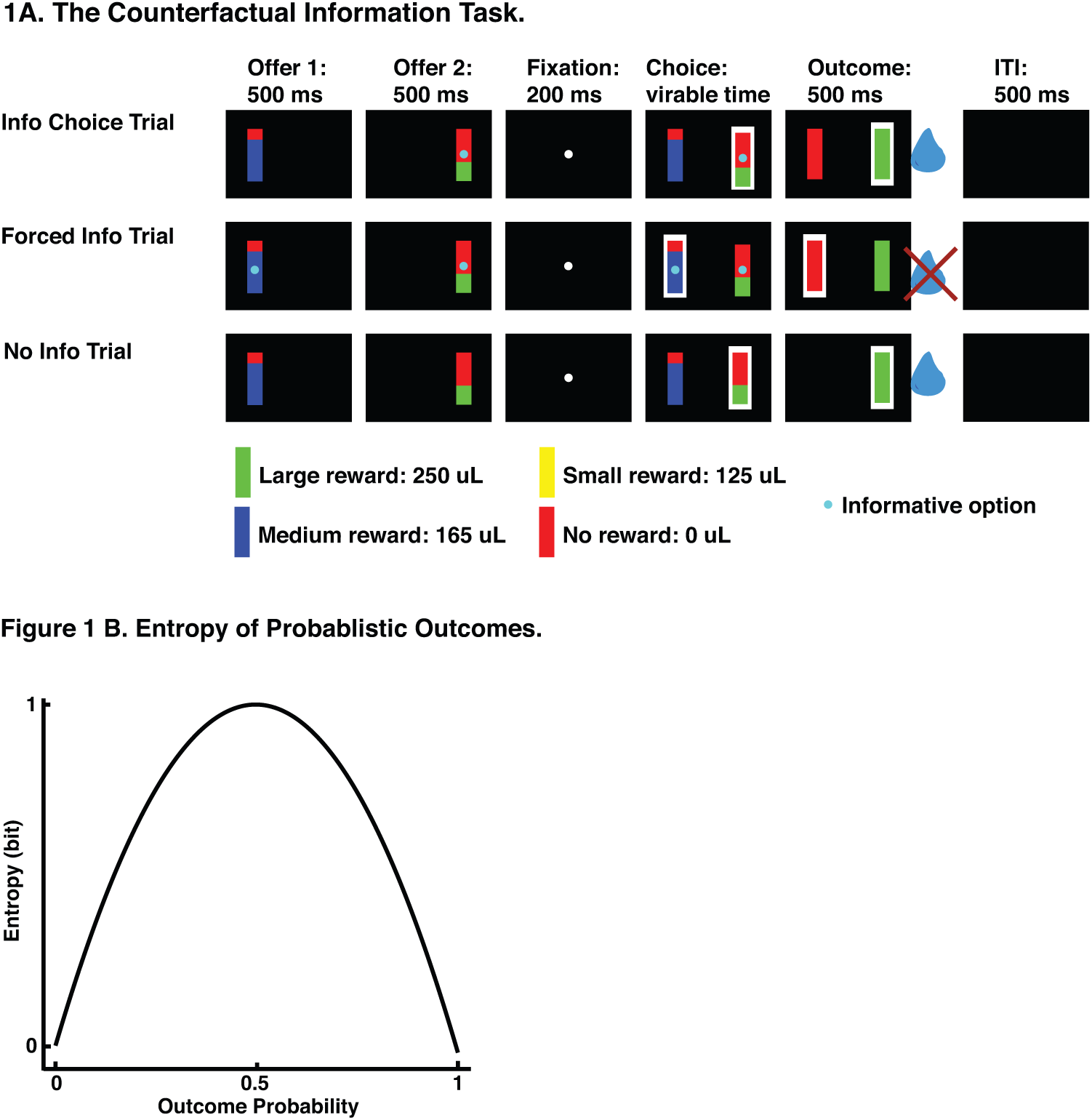
(A)The Counterfactual Information Task. Trial starts with serial presentation of offer 1 and offer 2, followed by a central fixation, choice, outcome, and an intertrial interval. The color of the offer indicates the size of the reward payoff and the height indicates the probability of receiving the payoff. The cyan dot at the center of the offers indicates the informativeness of an offer. Outcome is indicated following choice by filling the offer rectangle with the payoff color. Upon choice, payoff of the chosen gamble is delivered while the outcome is revealed. The informative option will lead to the reveal of both chosen and unchosen gamble outcomes. The non-informative option will lead to the reveal of only the chosen gamble outcome. On 50% of the trials, only one option was informative (*info choice* trials), on 25% of the trials, both options were informative (*forced info* trials), and on the rest, neither gamble was informative (*no info* trials). **(B) Entropy of Probabilistic Outcomes.** Entropy, information gained from revealing a gamble outcome, peaks at probability of 0.5, when the outcome is most uncertain, and decreases lawfully as probability moves closer to 0 or 1, when the outcome gets more certain.

We have previously used this general structure (without the informativeness manipulation) to probe macaques’ preferences for uncertainty and, through various controls, have demonstrated that they treat these as described gambles (Hayden, Heilbronner, & Platt, 2010; Heilbronner & Hayden, 2016). Critically, over the training period before data collection, subjects had ample opportunity to learn that the distribution of both actual outcome and counterfactual outcomes perfectly matched their probabilities. Monkeys’ behavior following training suggested that they understood the task. Most importantly, they chose the gamble with larger expected value 82.06% of the time (subject B: 81.68%; subject J: 82.69%). This proportion is larger than expected by chance (both subjects: X^2^=1865; P<0.001; subject B: X^2^=1133; P<0.001; subject J: X^2^= 729.83; P<0.001; chi-square test).

Monkeys preferred gambles that provided counterfactual information. To measure the effect of counterfactual information on choice, we used a multiple logistic regression model to fit the probability of choosing offer 1 as a function of four variables, the expected values and informativeness of the two offers. The probability of choosing offer 1 was positively predicted by expected value of offer 1 (B=0.0283, SE=0.0007, t-stat=38.94; P<0.001; **Figure 2 A-C**) and negatively predicted by that of offer 2 (B=-0.0270, SE=0.0007, t-stat=-37.50; P<0.001). By including informativeness in the same model, it competed with expected values to explain variance in behavior. We found that informativeness predicted choice above and beyond the effect of expected value. Specifically, the probability of choosing offer 1 was positively predicted by its informativeness (B=0.2633, SE=0.0616, t-stat=4.27; P<0.001; logistic regression) and negatively predicted by the informativeness of offer 2 (B=-0.2042, SE=0.0615, t-stat=-3.32; P=0.001; logistic regression). This result shows that monkeys were more likely to choose options with larger expected value and that provide counterfactual information.

**Figure 2.**
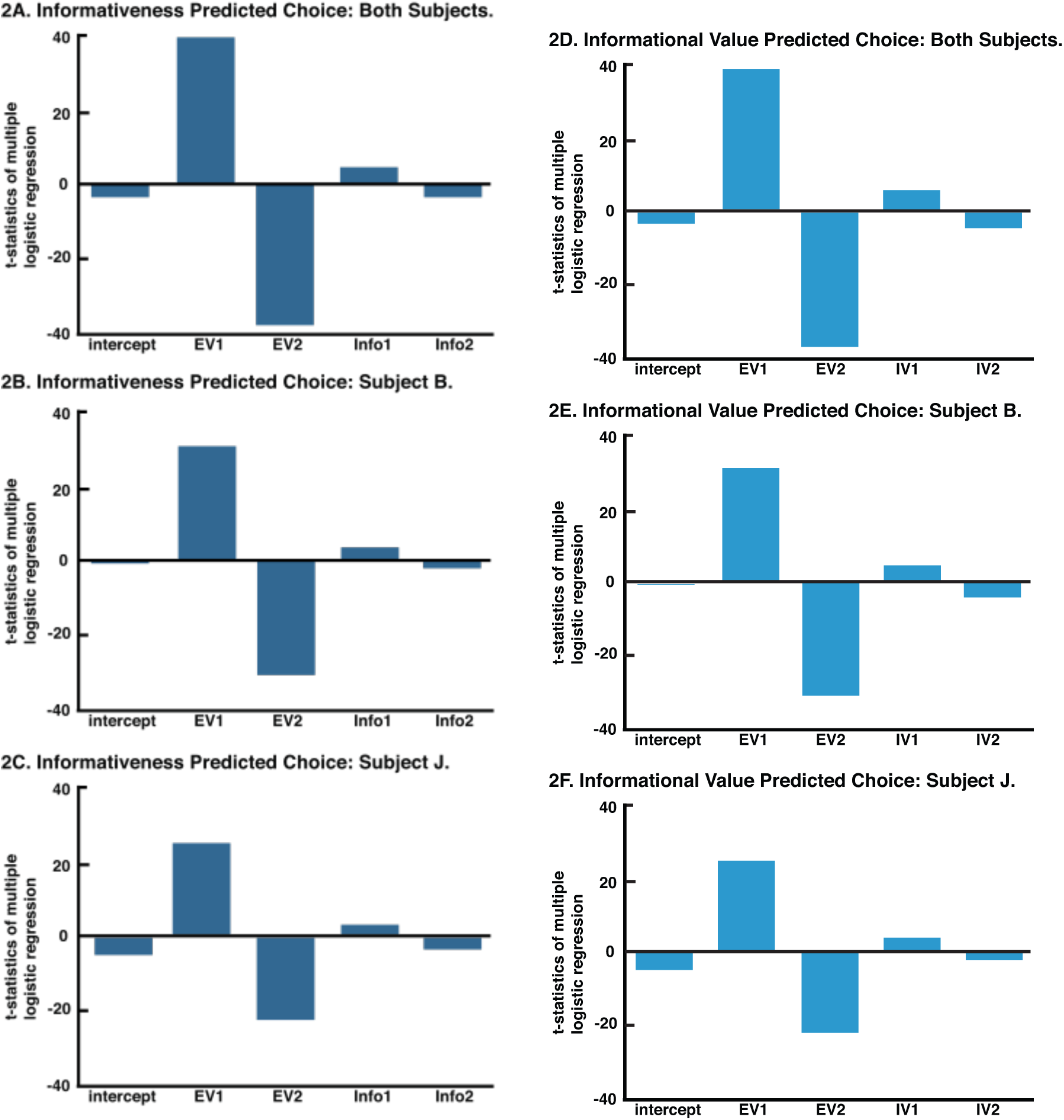
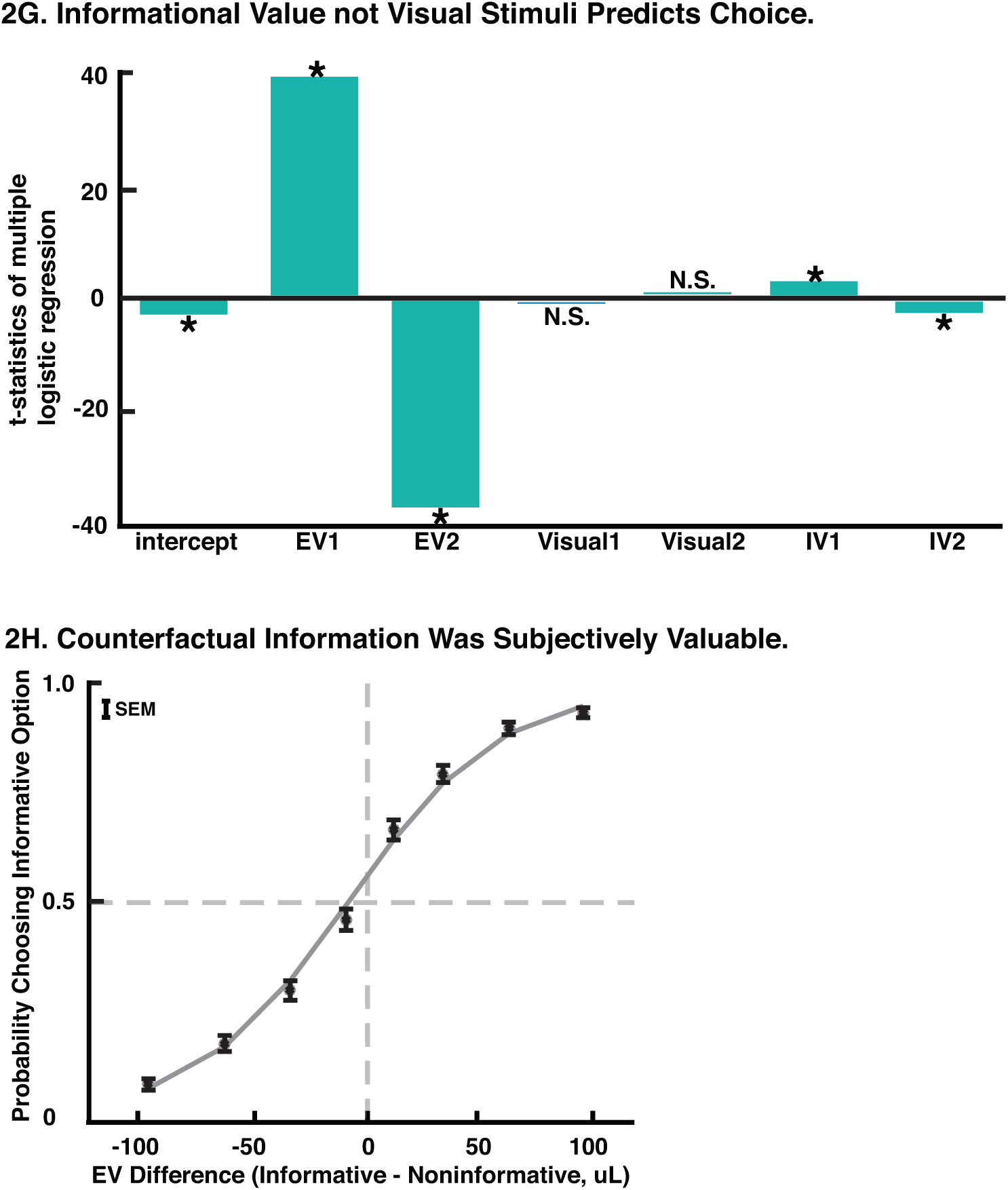
**(A-G)** X-axis:predictors included in the multiple logistic regression. Y-axis: t-statistics of each predictor. **(A-C)** Probability of choosing offer 1 as a function of expected values (EV) and informativeness (info) of offer 1 and offer 2, fitted for both subjects and each subject. **(D-F)** Probability of choosing offer 1 as a function of expected values (EV) and informational value (IV) of offer 1 and offer 2, fitted for both subjects and each subject. **(G)** Probability of choosing offer 1 as a function of expected values (EV), informativeness visual cues (visual), and informational value (IV) of offer 1 and offer 2. Asterisk: significant (p<0.05). N.S.: not significant. **(H)** Fitted logistic curve for probability of choosing informative option as a function of value difference (expected value of informative option minus expected value of non-informative option). The leftward shift shows the higher subjective value placed on informative options.

### Preference for counterfactual information scales with information

Observing the outcome of a gamble reduces uncertainty about the observed outcome and thus provides information in a formal sense (Cover & Thomas, 2006; MacKay, 2003; Shannon & Weaver, 2015). The amount of information, or entropy, provided by revealing a gamble outcome is not constant for all gamble probabilities. Instead, entropy, gained from revealing a gamble outcome peaks at probability of 0.5, when the outcome is most uncertain, and decreases lawfully as probability moves closer to 0 or 1, when the outcome gets more certain (**Figure 1 B**; Cover & Thomas, 2006; MacKay, 2003; Shannon & Weaver, 2015). Therefore, to satisfy our proposed definition of curiosity, subjects’ willingness-to-pay for counterfactual information should scale with the option’s entropy.

We defined the *informational value* (IV) of each gamble as the entropy of the chosen gamble when the chosen gamble was not informative and as the sum of entropy of both the chosen and unchosen gambles when the chosen gamble was informative (**Equation 2-4**; see **Methods**). We then used a multiple logistic regression model to fit the probability of choosing offer 1 as a function of expected values and informational values of the two offers. Note, critically, that we use the term informational value differently than informativeness, which is the binary variable we used in the previous section. Informational value refers to entropy, which is a continuous variable. Informativeness and informational value are orthogonal in our task because of the fully independently randomized probabilities for both options.

A logistic regression analysis that included informational value instead of informativeness revealed, first (not surprisingly), that monkeys preferred the higher value options. Specifically, probability of choosing offer 1 is positively predicted by the expected value of offer 1 (B=0.0281, SE=0.0007, t-stat=38.90; P<0.001; **Figure 2D-F**) and negatively predicted by that of offer 2 (B=-0.0268; SE= 0.0007; t-stat=-37.29; P<0.001). Second, and critically, the probability of choosing offer 1 was positively predicted by its informational value (B=0.3536; SE=0.0653; t-stat=5.42; P<0.001;) and negatively predicted the informational value of offer 2 (B=-0.2773; SE=0.0650; t-stat=-4.26; P<0.001). Thus, informational value explained a significant portion of variance in choice behavior, above and beyond that was explained by expected values. These results demonstrate that monkeys preferred options that provided higher entropy, and thus larger amount of information. Therefore, monkeys’ information-seeking tendency scaled with the amount of information provided.

### Monkeys prefer information, not the visual stimuli

One possible alternative explanation for monkeys’ preference is that they seek the options that have or that lead to *more* visual stimuli, which in this task are the informative options (Roper, 1999). To rule out this possibility, we conducted the following two analyses.

First, we entered expected value, informativeness (binary variable), and informational value (entropy associated with a choice) simultaneously into a multiple logistic regression (**Equation 6**; see **Methods**). Because informativeness was perfectly correlated with both presented and expected additional visual stimuli, this analysis allows the presence of additional visual stimuli and informational value to directly compete for explaining variance in choices. As before, we found that probability of choosing offer 1 is positively predicted by the expected value of offer 1 (B=0.0281, SE=0.0007, t-stat=38.74; P<0.001; **Figure 2G**) and negatively predicted by that of offer 2 (B=-0.0268; SE=0.0007; t-stat=-37.05; P<0.001;). Crucially, the probability of choosing offer 1 was not significantly predicted by the informativeness (i.e. presence of additional visual stimuli) of either offer 1 (B=-0.0248; SE=0.1190; t-stat=-0.21; P=0.835) or offer 2 (B=0.0312; SE=0.1193; t-stat=0.26; P=0.794). And yet, it was positively predicted by its informational value (B=0.3822; SE=0.1348; t-stat=2.84; P=0.005; logistic regression) of offer 1 and negatively predicted the informational value of offer 2 (B=-0.3095; SE=0.1343; t-stat=-2.31; P=0.021; logistic regression). These results indicate that the additional visual stimuli did not significantly account for variance in the behavior. It is the Shannon information, not the additional visual stimuli associated with informative options *per se* that drives preference.

Second, if monkeys’ preference for informative gambles truly reflected their tendency to seek more information, then using informational value, instead of informativeness, in the multiple logistic regression would yield a better fit to the choice behavior. Akaike information criterion (AIC) is one of the common measurements for formal model comparison. We found the model incorporating entropy resulted in a smaller AIC score (AIC= 6571.97) than the previously described model (AIC= 6580.22), indicating a better fit to the data. AIC weight estimates the relative likelihood of a particular model among all the candidate models, and thus provides a quantitative measure of how much better a model is to the alternative(s) (Burnham & Anderson, 2010). Comparing these two AIC values, the model incorporating entropy resulted in an AIC weight of 98.41%, which means that this model is 98.41% more likely to be the one that resembles the true data-generating model, and thus better describing the choice behavior (this percent roughly translates to P=0.016 in traditional significance test; **Equation 7**; see **Methods**). These results further confirm that monkeys’ preferences for the informative gamble was driven by its informational value, i.e. entropy, rather than driven by potential confounds, such as total visual stimuli on the screen.

### Monkeys do not use counterfactual information to update choice strategy

Although counterfactual information in our task provided no strategic benefit due to trial-to-trial independency, we wondered whether monkeys nonetheless acted as if it did; if so they might have adjusted their strategy after receiving counterfactual information (Hayden et al., 2009). We thus examined changes in preference resulting from counterfactual information. Choice accuracy (likelihood of choosing the option with the greater expected value) did not measurably change after receiving counterfactual outcome information. Specifically, it was 81.48% (n= 4227) when the counterfactual outcome was revealed and 82.68% (n= 3914) when it was not (X^2^=1.99; P=0.158; chi-square test).

The valence of counterfactual information also did not measurably affect subsequent choices. Counterfactual information could potentially lead to either a *good news* condition (chosen gamble win and unchosen gamble loss, or, chosen win > unchosen win), or a *bad news* condition (chosen gamble loss and unchosen gamble win, or, chosen loss > unchosen loss). We found no difference in subsequent choice accuracy following a good news condition (82.57%) versus a bad news condition (81.02%; X2=2.24; P=0.134; chi-square test). Moreover, a bad news condition (possibly leading to a regret-like state) did not motivate choice of the unchosen side (X^2^=0.05; P=0.818; chi-square test) or original position (X^2^=0.07; P=0.789; chi-square test) relative to a good news condition, and thus ruling out win-stay-lose-shift strategy based on counter-factual information. We also found no difference in subsequent information-seeking tendency following a good new condition versus a bad news condition (53.46% versus 52.45%; X^2^=0.26; P=0.609; chi-square test). These results show that counterfactual information did not lead to measurable choice strategy shifts and argue against the possibility that our effects reflect erroneous belief by our subjects about the meaning of the signals they see.

### Measuring the value of curiosity for counterfactual information

To quantify the subjective value placed on counterfactual information, we generated psychometric curves showing probability of choosing the informative gamble as a function of expected value difference between informative and non-informative gambles (**Figure 2H**). This curve was shifted to the left, indicating that monkeys sacrificed water reward for information. On average, they sacrificed 6.40 uL of water reward (subject B: 6.41 uL; subject J: 6.37 uL) relative to a pure reward-maximizing strategy (t-stat=4.87; P<0.001; t-test), to gain 0.014 bits (subject B: 0.0147 bits; subject J: 0.0135 bits) more information. This water payment for information is 5.32% the size of the average reward obtained per trial. Thus, monkeys sacrificed a small but significant amount of primary reward to satisfy their curiosity about the counterfactual outcomes.

## DISCUSSION

We find that when choosing between risky options, macaques prefer gambles that promise information about what would have occurred had they chosen differently. This information, known as counterfactual information, has no direct or indirect benefits and provides no information about the statistics of the current task environment. Indeed, we see no measurable effect of counterfactual outcomes on strategic adjustments. As such our task satisfies the three strict criteria we propose for human-like curiosity in non-human animals: (1) a willingness to pay for information that (2) gives no strategic benefit and (3) the willingness to pay scales with amount of information available.

Many non-human animals will explore new environments or stimuli (Kidd & Hayden, 2015). Although exploratory behaviors may reflect curiosity, it is hard to ascertain whether they reflect a drive for information *per se*. For this reason, many scholars favor more controlled paradigms. Many of these use observing behavior – a preference to reduce uncertainty about stochastic upcoming rewards. Observing behavior has been shown in numerous species; however, previous results have not established that the observing behavior scales with entropy (Badia, Ryan, & Harsh, 1981; R. Blanchard, 1975; Blanchard et al., 2015a; Bromberg-Martin et al., 2010; Dinsmoor, Mulvaney, & Jwaideh, 1981; Lieberman, 1972; Prokasy, 1956; Purdy & Peel, 1988; Roper, 1999; Steiner, 1967; Wilton & Clements, 1971; Wyckoff, 1952). Due to this lack of support from formal information theory, observing behavior suffer from a potential confound with Pavlovian learning processes (Beierholm & Dayan, 2010; Dayan et al., 2006; Roper, 1999).

In this light, another potential limitation is the possibility that seemingly information-seeking behaviors could have been a byproduct of other processes. One such process is when information gained is a byproduct as animals explore and manipulate the environment to acquire reward (Emery & Clayton, 2004; Thorndike, 2017). Another example is that information seeking leads to learning of the structure of a task and thus to strategic benefits (Daw et al., 2006; Doll, Duncan, Simon, Shohamy, & Daw, 2015; Noonan et al., 2010; Walton et al., 2010). Yet another case is that information could be misconceived as influencing the likelihood of reward (Heil-bronner & Hayden, 2013; Vasconcelos et al., 2015). In all cases, information is acquired not for the gain of information but for the gain of other perceived instrumental benefits.

In the current task, we provided subjects with the opportunity to actively choose to reveal counterfactual outcomes, even at the cost of primary rewards. Due to the cross-trial independent design, counterfactual information provides neither immediate or future rewards, nor actual or perceived strategic benefits. The lack of strategic adjustment after receiving counterfactual information in our subjects confirmed the effectiveness of this manipulation. Importantly, animals’ willingness to pay scaled with the amount of information. This result reflects the idea that curiosity must necessarily be a drive for information. That is, to use Loewenstein’s term, it resolves an *information gap* between current knowledge and potential knowledge (Golman & Loewenstein, 2015; 2016; Gruber et al., 2014; Loewenstein, 1994). Therefore, our results bridged the theoretical and empirical findings with formal information theory to show human-like curiosity in non-human primates.

Our formal criteria for curiosity include a requirement that amount of curiosity scales with amount of information available. This criterion is important because it explicitly links ostensible curiosity behavior to an information gap per se. It also provides a ready control for reinforcement processes, such as engagement to cues and tracking of conditioned reinforcers. Note that, formally speaking, we only require a positive relationship between entropy and willingness to pay along a subset of possible values. Our criteria allow future research the possibility for more complex relationship along a full range. For example, it is likely that demand form information eventually saturates and even decreases as entropy rises (Kidd, Piantadosi, & Aslin, 2012; 2014).

These results have direct implications for neuroscientific research. The goal of understanding curiosity is important for its role in driving choice and learning in uncertain environments (Kidd & Hayden, 2015). We offer a well-controlled paradigm for isolating curiosity from potential confounds separating it from seemingly similar factors. One neuroscientific problem that could benefit from our result is how curiosity influences learning and reward processes. Particularly, understanding how prospective and received values of information outcome and primary reward outcome are represented, and how curiosity or value of information influences learning and update of model-free and model-based processes, across prefrontal, striatal, and ventral tegmental areas, would start to reveal how curiosity could serve as an intrinsic drive to shape an organism’s learning and decision-making (Chang, Gardner, Di Tillio, & Schoenbaum, 2017; Farovik et al., 2015; Gershman & Schoenbaum, 2017; Howard & Kahnt, 2017; Kahnt, Heinzle, Park, & Haynes, 2010; Langdon, Sharpe, Schoenbaum, & Niv, 2018; Sadacca et al., 2018; Stalnaker, Liu, Takahashi, & Schoenbaum, 2018; Takahashi et al., 2017; Wang & Hayden, 2017).

## METHODS

### General Methods

All animal procedures were approved by the University of Rochester Animal Care and Use Committee and were conducted in compliance with the Public Health Service’s Guide for the Care and Use of Animals. Two male rhesus macaques (*Macaca mulatta*), aged 9-10 years and weighting 8.0-9.9 kg served as subjects. Both subjects had extensive previous experience in risky decision-making tasks. Subjects had full access to food (LabDiet 5045, ad libitum) while in their home cages. Subjects received at minimum 20 mL per kg of water per day, although in practice they received close to double this amount in the lab as a result of our experiments. No subjects were sacrificed as a result of these experiments, nor were they physically harmed.

Visual stimuli were colored rectangles on a computer monitor (see Figure 1). Stimuli were controlled by Matlab with Psychtoolbox and Eyelink Toolbox. A solenoid valve controlled the delivery duration of fluid rewards. Eye positions were sampled at 1,000 Hz by an infrared eye-monitoring camera system (SR Research, Osgoode, ON, Canada). A small mount was used to facilitate maintenance of head position during performance.

Subjects had never previously been exposed to decision-making tasks in which counter-factual information was available. Previous training history for these subjects included two types of foraging tasks (T. C. Blanchard & Hayden, 2015; T. C. Blanchard, Strait, & Hayden, 2015b), intertemporal choice tasks (Hayden, 2016), two types of gambling tasks (Azab & Hayden, 2017; Strait, Blanchard, & Hayden, 2014), and two types of reward-based decision tasks(Sleezer, Castagno, & Hayden, 2016; Wang & Hayden, 2017).

### The *Counterfactual Information* Task

Two subjects (B and J) performed a novel task designed to measure preference for counterfactual information (**Figure 1A**). On each trial subjects chose between two randomly selected gambles, presented asynchronously on the left and the right side of the screen. Gambles were represented as rectangular visual stimuli and differed in three dimensions: payoff, probability, and *informativeness*. Payoff came in three sizes, small (125 microliters), medium (165 microliters), and large (250 microliters), each corresponding to a yellow, blue, and green portion of the rectangle. Probabilities were randomly drawn from a uniform distribution between 0 and 1 (step size of 0.01). The height of a portion of the rectangle with a payoff color indicated probability of winning the gamble and the height of the red portion indicated the probability of losing the gamble (that is, of receiving no reward for that trial). Informativeness of a gamble was indicated by a cyan dot on the center of the rectangle for an informative option and the lack of a cyan dot for a non-informative one. The informative option promised valid information about the payoff that would have occurred had the alternative option been chosen. On 50% of the trials, only one option was informative, on 25% of trials, both gambles were informative, and on the remaining 25% of trials, neither gamble was informative. Probability, payoff, and informativeness were independently randomized on each trial.

Each trial started with the appearance of offer 1 (500 ms) followed by a blank 500 ms delay. Offer 1 position was randomized for each trial. Then offer 2 appeared on the other side of the screen (500 ms) followed by another 500 ms delay. After a 200 ms fixation, both gambles appeared on the screen and subjects chose the preferred option by shifting gaze to it and maintaining that gaze for 200 ms. Subsequently, if an informative option was chosen, gamble results for both offers were resolved. If a non-informative option was chosen, gamble results for only the chosen offer was resolved. Resolution of a gamble involved filling the gamble rectangle with the payoff color, while delivering a water reward, if the gamble result was win, or filling the gamble rectangle with red color, while delivering no reward, if otherwise. This outcome epoch lasted for 800 ms and was followed by a 1000 ms inter-trial interval (ITI) and then the start of next trial.

## Statistical Methods

All choices were counted as correct when subjects selected an option with expected value greater than or equal to the non-chosen alternative. Chi-square test was conducted using R. Subjects’ choice behavior was fitted using a multiple logistic regression model and was conducted using MATLAB (Mathworks). T-test was also carried out with MATLAB.

A logistic regression was fitted to choice to assess whether subjects preferred informative option, above and beyond the effect of expected value:

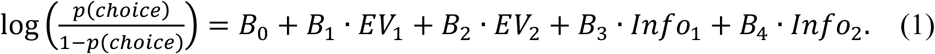

EV stands for expected value, which is the product of reward magnitude and reward probability. Info is 1 when choice of an option leads to resolution of both chosen and unchosen gambles and is 0 when it leads to the resolution of only the chosen gamble.

To quantify the amount of information provided when a gamble outcome is resolved, we calculated the “information value” (IV) of each option. The IV is the uncertainty about the possible outcomes that will be eliminated by either observing the outcome or receiving information:

the Shannon entropy (H) of the offer:

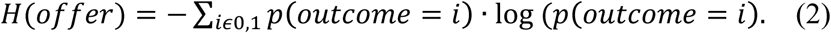

P is the reward probability associated with a gamble option.

When the non-informative offer is chosen, the informational value (IV) of this choice is:

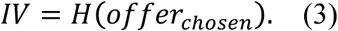

When the informative offer is chosen, the informational value (IV) of this choice is:

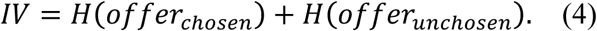

A separate logistic regression was fitted to choice to assess whether subjects preferred choice with higher informational value, above and beyond the effect of expected value:

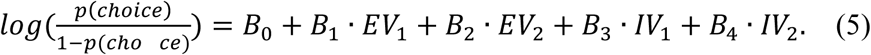

A third logistic regression was fitted to allow for EV, the additional visual stimuli (Visual) that came with informativeness options, and informational value to compete to explain variance in choice:

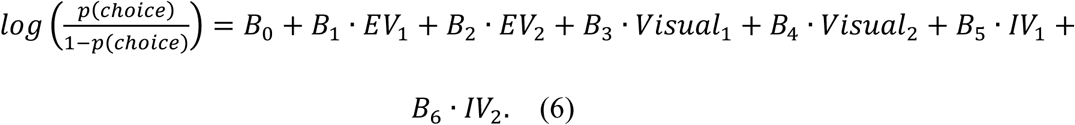

For model comparison, AIC weights was calculated as following:

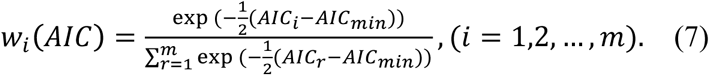

W_*i*_ is probability of a model M_*i*_being the one, among all *m* candidate models that is most close to the true data-generating model (Burnham & Anderson, 2010).

## Data Availability

The datasets generated during the current study are available on the Hayden lab website, http://www.haydenlab.com/, or from the authors on reasonable request. The analyses code generated during the current study is available from the corresponding author on reasonable request.

## Author Contributions

B.Y.H and M.Z.W designed the experiment, M.Z.W conducted the experiment and analyzed the data, B.Y.H and M.Z.W wrote the paper.

## Acknowledgements

We thank Shannon Cahalan, Marcelina Martynek, and Michelle Ficalora for help with data collection and Becket Ebitz for useful comments on the manuscript. This research was supported by a grant to B.Y.H from NIH R01 DA037229.

